# TAAC - TMS Adaptable Auditory Control: a universal tool to mask TMS click

**DOI:** 10.1101/2021.09.08.459439

**Authors:** S. Russo, S. Sarasso, G.E. Puglisi, D. Dal Palù, A. Pigorini, S. Casarotto, S. D’Ambrosio, A. Astolfi, M. Massimini, M. Rosanova, M. Fecchio

## Abstract

**Background:** Coupling transcranial magnetic stimulation with electroencephalography (TMS-EEG) allows recording the EEG response to a direct, non-invasive cortical perturbation. However, obtaining a genuine TMS-evoked EEG potential requires controlling for several confounds, among which a main source is represented by the auditory evoked potentials (AEPs) associated to the TMS discharge noise (TMS *click*). This contaminating factor can be in principle prevented by playing a masking noise through earphones.

**New method:** Here we release TMS Adaptable Auditory Control (TAAC), a highly flexible, open-source, Matlab^®^-based interface that generates in real-time customized masking noises. TAAC creates noises starting from the stimulator-specific TMS *click* and tailors them to fit the individual, subject-specific *click* perception by mixing and manipulating the standard noises in both time and frequency domains.

**Results:** We showed that TAAC allows us to provide standard as well as customized noises able to effectively and safely mask the TMS *click*.

**Comparison with existing methods:** Here, we showcased two customized noises by comparing them to two standard noises previously used in the TMS literature (i.e., a white noise and a noise generated from the stimulator-specific TMS *click* only). For each, we quantified the Sound Pressure Level (SPL; measured by a Head and Torso Simulator - HATS) required to mask the TMS *click* in a population of 20 healthy subjects. Both customized noises were effective at safe (according to OSHA and NIOSH safety guidelines), lower SPLs with respect to standard noises.

**Conclusions:** At odds with previous methods, TAAC allows creating effective and safe masking noises specifically tailored on each TMS device and subject. The combination of TAAC with tools for the real-time visualization of TEPs can help control the influence of auditory confounds also in non-compliant patients. Finally, TAAC is a highly flexible and open-source tool, so it can be further extended to meet different experimental requirements.

## 1. Introduction

The TMS evoked EEG potential (TEP) reflects the activation of cortical neurons underneath the TMS coil to the extent that confounding factors are effectively controlled for (Belardinelli et al., 2019; Conde et al., 2019). Specifically, while electromagnetic artifacts can be controlled by optimizing the experimental setup, TEPs can be still contaminated by undesired biological activations – some of which can be identified and minimized in real-time such as muscle artifacts (Casarotto et al., 2021). In addition to these more easy-to-control confounds, TMS pulses vibrate the coil, thus generating a tapping sensation on the scalp and a loud noise denoted as TMS *click* that may result in sensory-related activations(Conde et al., 2019; Miniussi and Thut, 2010; Nikouline et al., 1999; Rogasch et al., 2014). For this reason, the TMS-EEG community recently stressed the importance of avoiding such confounds to obtain a genuine TEP (Belardinelli et al., 2019).

Recently, Rocchi and colleagues aimed at disentangling the role of these confounding factors and reported that AEPs represent the main source of undesired peripheral co-activation to TEPs (Rocchi et al., 2021). To rule out AEP from TEP, two main approaches have been employed: one approach relies on mathematically removing the AEP from the TEP (Miniussi and Thut, 2010; Nikouline et al., 1999; Rogasch et al., 2014), assuming a linear summation between them; by contrast, another approach prevents the AEP by masking the TMS *click* through the continuous delivery of a background noise to the subject. The second approach aims to effectively mask the perception of the TMS *click* (ter Braack et al., 2015) by playing either a white noise (Paus et al., 2001) or a noise adapted to the TMS *click* (Massimini et al., 2005) during TEP acquisition. Although arguably more effective, this second approach is limited by the lack of a general procedure for the noise generation, and by the risk of delivering excessively loud noises reported in previous TMS-EEG studies (Conde et al., 2019; Ozdemir et al., 2021; ter Braack et al., 2015).

To overcome these limitations, here we present TMS Adaptable Auditory Control (TAAC), a software that allows generating in real-time a masking noise adapted on the device-specific TMS *click* and tailored to abolish the subject-specific perception of the TMS *click*. While previous masking noises could accommodate subjects’ perception only by increasing the loudness, TAAC designs customized noises that accommodate subjects’ perception by optimizing the noise also in the time and frequency characteristics. To showcase the benefit of using TAAC, we performed psychophysical measures (quantified by accurate Sound Pressure Level – SPL measurements) to systematically compare the customized masking sound with the standard masking methods represented by a white noise and a noise adapted on the TMS *click* only.

## 2. Materials and methods

We designed TAAC (Fig. 1A) to optimally minimize the impact of auditory confounds during TMS-EEG experiments. TAAC is a Matlab^®^-based (The Mathworks) open-source (freely available at www.github.com/iTCf/TAAC) Graphic User Interface that allows generating a masking noise based on the recording of the original *click* of the stimulator and coil (i.e., device-specificity) and customized through subject-tailored distortions (i.e., subject-specificity).

**Figure 1:**
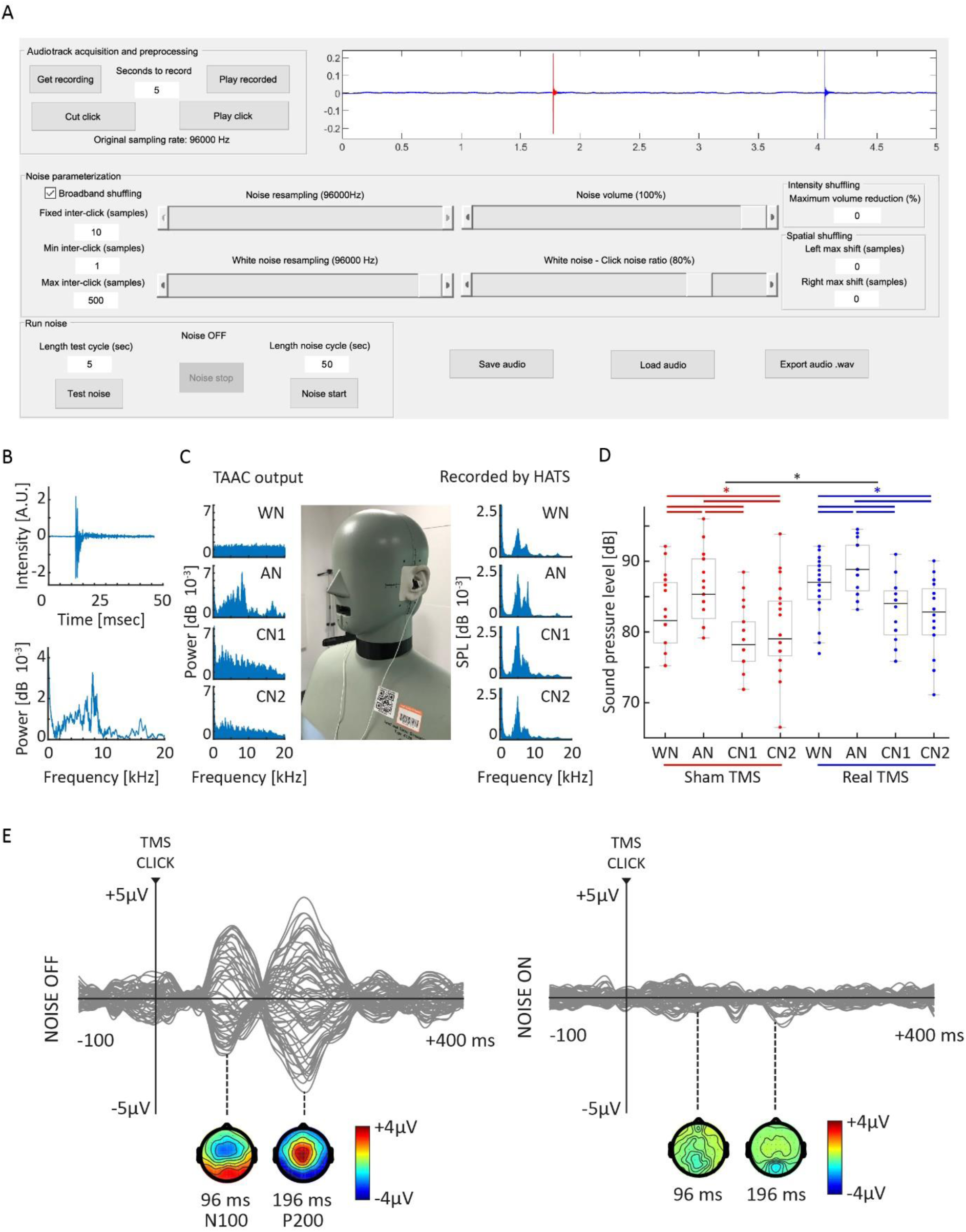
TAAC, psychophysical and electrophysiological test. **Panel A**: TAAC Graphic User Interface.; **Panel B**: Characterization of the selected *click* in time (top) and frequency domains (bottom); **Panel C**: Left: Spectra of the masking noises generated using TAAC. Middle: Photo of the setup employed to quantify SPL. Right: Spectra of the masking noises recorded through the HATS mannequin; **Panel D**: Boxplots illustrate the minimum SPL required to mask TMS *click* for each condition (red: sham TMS; blue: Real TMS) and masking noise. Each subject is depicted as a dot. Top lines show significance values of the post-hoc statistics (* = p<0.05); **Panel E**: Butterfly plot of the EEG response evoked by TMS *clicks* delivered without masking noise (left) and during noise administering (CN1, right). Instantaneous voltage topographies show the localization of the N100 and P200 components typical of the AEP (left) which are completely abolished during masking noise administration (right).

### 2.1. Description of the tool

To generate the customized masking noise using TAAC, TMS *click* is acquired through the *Audiotrack acquisition and preprocessing* panel, played through the *Run noise* panel, and customized through the *Noise parameterization* panel. Below is a brief description of these panels of the TAAC user interface. TAAC requires the Signal Processing toolbox and the Communications System Toolbox.

#### 2.1.1. Step 1: Audiotrack acquisition and preprocessing

The TMS *click* can be recorded with an omnidirectional microphone (without frequency distortions) placed 25 cm laterally to the TMS coil (Koponen et al., 2020) while delivering a few TMS pulses in a room with low noise and reverberation. The *Get recording* button enables the audio recording (sampling rate: 96.000 Hz, which is the maximum allowed by Matlab) of the duration specified in the *Seconds to record* box. Subsequently, the *Cut click* button allows for cropping the recording of a specific TMS *click* by isolating the time-window including the TMS *click* (temporal and spectral characterization of the cropped *click* in Fig. 1B). To verify the quality of the audio tracks, the *Play recorded* and *Play click* buttons play the entire soundtrack and the cropped TMS *click,* respectively.

#### 2.1.2. Step 2: Run noise

The noise can be played through the *Test noise* and the *Noise start* buttons. While *Noise start* is meant to be used in real experimental sessions, *Test Noise* is meant to be employed before the experiment, to test and iteratively adjust the noise parameters.

Specifically, the *Test noise* button creates a noise of a certain length (as specified in the *Length test cycle* box) that is iteratively played, so that the user can real-time manipulate some of the noise parameters. Hence, the noise iteratively changes according to the modifications in the parameters.

Instead, the *Noise start* button creates a noise of a certain length (as specified in the *Length noise cycle* box) and iteratively plays it while all the parameters are locked.

#### 2.1.3. Step 3: Noise parameterization

Fine masking of the TMS *click* can be achieved by tuning the features of the noise on the perception of the TMS *click* of each subject by adjusting the noise parameters (See supplementary methods) based on the real-time subject’s feedback and/or on the presence of the AEP (Casarotto et al., 2021). In this manner, TAAC can be useful to mask TMS *clicks* also in non-compliant subjects.

The customized noise delivered to the subject is composed of two independent parts: the *click*-based noise (CBN) and the white-based noise (WBN). CBN is derived from the recorded TMS *click* whereas WBN is randomly generated.

CBN originates from the TMS *click* recorded and cropped by the user (see Step 2). Specifically, TAAC creates a sequence of TMS *clicks* concatenated with pseudo-random shifts. These shifts (measured in samples) are taken from a uniform distribution (Minimum: *Min inter-click* box; Maximum: *Max inter-click* box) added to a fixed inter-*click* interval (i.e., *Fixed inter-click* box).

According to the subject’s feedback, by moving the *Noise frequency* bar, CBN can be resampled towards a higher or lower frequency, thus affecting the recorded *click* pitch (i.e., towards a higher or lower central frequency). The *BroadBand shuffling* checkbox automatically designs a pre-customized CBN that encompasses the original TMS *click* mixed with several distorted versions of the *click* itself - each one resampled with a different frequency either above or below the original one. This approach excludes the CBN frequency tuning (i.e., *Noise frequency* bar), reducing the degrees of freedom of the noise parameterization and the time required to parameterize the masking noise.

By sliding the *White noise frequency* bar, WBN is resampled minimizing the high frequencies with respect to the lower frequencies. To balance the proportion of CBN and WBN, the *White noise – Click noise ratio* bar establishes the percentage of WBN and CBN in the final noise. Then, the *Noise volume* bar modulates the intensity of the noise independently from the computer volume.

TAAC also provides supplementary features. An example already implemented in the toolbox is provided by the *Intensity Shuffling Box*. In this case, the intensity of each *click* of the CBN can be randomly adjusted over time (through a uniform distribution) ranging from 0% to the percentage specified in the *Maximum volume reduction* box. Another example is provided by the *Spatial shuffling* box. In this case, the virtual location of the TMS *click* source along the transversal axis can be also modulated and randomized by introducing a time delay between the ears (as specified by the user in the *Left max shift* and *Right max shift* box). Although these features were not tested in the current paper – they were set to 0 - they could further improve the quality of the masking noise and possibly reduce the sound pressure level in particular conditions – such as extremely lateralized TMS stimulations and randomized pulse intensities.

Finally, the whole set of parameters can be saved (i.e., *Save audio* button) and reloaded (i.e., *Load audio* button), guaranteeing a high reproducibility of the masking noise across repeated TMS stimulation sessions.

### 2.2. Noise administering

An optimal masking of the TMS *click* can be achieved by delivering the noise through in-ear earphones (Conde et al., 2019; Fecchio et al., 2017; Rocchi et al., 2021) that help to acoustically isolate the subject from the environment. The noise can be played directly from Matlab, or it can be exported to .wav format through the *Export audio .wav* button. In this manner, once the masking noise has been customized, it can be played through other devices (e.g., smartphones).

### 2.3. Testing the tool

To showcase the advantage provided by TAAC, we measured the minimum SPL required to mask the TMS *click* by applying four different types of noise administered through in-ear earphones. In particular, we compared the noises previously used in literature, i.e. white noise (WN) and adapted noise (AN: noise adapted on the TMS *click* as in Massimini et al., 2005), with two customized noises (CN1 and CN2) generated through the *Broadband shuffling* function (noise adapted on the TMS *click,* customized through a broadband shuffling, and mixed with white noise with two different parameterizations – for details see Table S1). We performed these measurements in a group of 20 healthy subjects (F=10; Age range 24-60 years). Also, we anecdotally confirmed that the AEP can be effectively abolished by using TAAC in one subject. All the participants provided written informed consent. Experiments were approved by Ethics Committee Milano Area A. Exclusion criteria included history of CNS active drugs, of abuse of any drug, and of neurological or psychiatric disease.

#### 2.3.1. Sound pressure level measurement

SPLs were measured in the Audio Space Lab of the Department of Energy of Politecnico di Torino, which is characterized by low reverberation (T30<0.3 s at mid-frequencies) and low background noise (LAeq<37dB). A Head and Torso Simulator (HaTS, model 4128 by Bruel & Kjaer) was equipped with the same in-ear earphones (model JVC HA-FX8) used with real subjects, connected to the personal computer where the TAAC tool was installed. A digital analyzer (model 2900 by Larson Davis) was used to read SPL values for all noises and volumes. As all the tested noises had a stationary component in time, they were acquired and evaluated as A-weighted running SPLs to account for the filtering of the human ear (Fig. 1C).

#### 2.3.2. Psychophysical test of noise efficacy and safety

For each masking noise (WN, AN, CN1, and CN2), we assessed the minimum SPLs at which the 20 subjects involved in the study did not report any perception of the TMS *click*. The TMS *clicks* used to test the performances of TAAC were generated by delivering single TMS pulses (Focal figure-of-eight coil, biphasic pulse shape, diameter 50/70 mm, driven by an NBS4 stimulation unit, Nexstim Ltd., Finland) at 70% of the maximum stimulator output (approximated maximum stimulator output from Fecchio et al., 2017) in a Real TMS condition (coil placed 2 cm posteriorly to the vertex to the scalp; induced electric field oriented along the anteroposterior axis) and in a sham TMS condition (i.e., coil placed at the vertex of the subject’s scalp and tilted at 90° on the sagittal plane touching the scalp).

For each masking noise, while delivering TMS pulses (at a jittered inter-pulse interval between 2000-2300 ms), we progressively increased the volume intensity until the subject was not able to detect any TMS *click* (10 consecutive undetected pulses). Then, for each noise type and condition, we assessed the volume intensity at which each subject did not report any perception of the TMS *click*. To avoid entrainment and sorting effects, the order of the conditions and noise types was randomized for each subject. From each value of the computer volume and for each noise type, we extrapolated the corresponding SPL value that was able to mask the TMS *click*. Finally, for the SPL obtained in each subject, we assessed the maximum exposure time according to the safety tables of the US National Institute for Occupational Safety and Health (NIOSH) and the US Occupational Safety and Health Administration (OSHA).

#### 2.3.3. Electrophysiological proof-of-concept for noise efficacy

As a proof of concept for the efficacy of the proposed masking noise in abolishing the AEP during TMS-EEG experiments, we performed a sham TMS-EEG recording session in subject 4. This experiment also serves as an example of how TAAC can be associated with a real-time interface for TMS-EEG (Casarotto et al., 2021) to ensure the lack of auditory confounds. Specifically, we acquired one EEG session while delivering sham TMS without noise masking (as null control) and one EEG session during the supply of customized noise generated with TAAC (CN1 administered at the minimum SPL at which the subject did not report any TMS *click*).

During each TMS-EEG session, we delivered 120 sham TMS pulses at 70% of the maximum intensity and these data were preprocessed using custom-made Matlab^®^-based scripts (see supplementary methods).

#### 2.3.4. Statistical analysis

SPL values were compared at the group level between conditions and among noise types by 2-ways Analysis of Variance – ANOVA followed by post-hoc pairwise t-test (Holm-Bonferroni correction for multiple comparison) using R 4.1.0.

## 3. Results

To illustrate the advantage of using TAAC with respect to existing masking noises, we compared the minimum SPL whereby TMS *click* was effectively masked by WN, AN, CN1 and CN2. The two customized noises (CN1 and CN2) generated through TAAC effectively masked the TMS *click* at a lower SPL with respect to WN and AN. Specifically, the median value across subjects of the minimum SPL required to mask the TMS *click* with WN, AN, CN1 and CN2 noises was 81.6, 85.4, 78.3, and 79.0 dB, respectively in the sham TMS condition and 87.1, 89.0, 84.1, and 82.9 dB, respectively in the Real TMS Condition.

ANOVA revealed a significant main effect of both noise type and condition with no interactions (interaction, *F*_(3,57)_=2.097, *p*=0.11; noise, *F*_(3,57)_=73.667, *p*<0.001; condition, *F*_(1,19)_=22.181, *p*<0.001). Post-hoc pairwise t-tests (Holm-Bonferroni corrected) revealed that all comparisons between conditions and between noise types were significant (p<0.05), except for CN1 vs CN2. Table S1 reports the single-subject equivalent SPL values at which each noise was able to mask the TMS *click* in each condition.

Both during Real and Sham TMS conditions, the minimum SPL value between the two customized noises CN1 and CN2 was always lower than the SPL value of both WN and AN in all subjects. Specifically, in the sham TMS condition, the customized noises (minimum value between CN1 and CN2) provided a median reduction of 4.1 dB with respect to WN and of 7.5 dB with respect to AN (reduced in 100% of the subjects); in the Real TMS condition, the customized noises provided a median reduction of 5.6 dB with respect to WN and a median reduction of 7.8 dB with respect to AN (reduced in 100% of the subjects). In terms of sound intensity (W/m^2^), these SPL reductions are equivalent to a reduction of 61%, 82%, 72%, and 83%, respectively.

We related the SPLs of the Real TMS condition to two international scales for exposure safety (i.e., NIOSH and OSHA), finding that WN and AN can be administered for median latencies adequate for TMS-EEG experiments (WN NIOSH: 5 hours; WN OSHA: 12 hours; AN NIOSH: 3 hours; AN OSHA: 9 hours;), while customized noises can be administered for even longer times (CN1 NIOSH: 10 hours; CN1 OSHA: 18 hours; CN2 NIOSH: 13 hours; CN2 OSHA: 21 hours;).

In one subject, we additionally recorded the EEG response to a sham TMS session without masking noise and with masking noise (CN1). At the end of the measurement without masking noise, the subject reported to be able to hear all the delivered TMS *clicks,* as also documented by the presence of a typical AEP in the EEG response (Fig 1E, left side). At the end of the measurement with masking noise (CN1), the subject reported not being able to hear TMS *clicks,* as also documented by the absence of any evoked EEG response (Fig 1E, right side).

## 4. Discussion

TMS-EEG experiments performed without controlling for the auditory component result in TEPs that are contaminated by auditory potentials (Massimini et al., 2005; Nikouline et al., 1999; Rocchi et al., 2021; Rogasch et al., 2014; ter Braack et al., 2015). Controlling for the auditory confounding can be challenging, so that merely increasing the loudness of the masking noises has proved to be not always effective in previous TMS-EEG studies (Conde et al., 2019; Ozdemir et al., 2021; ter Braack et al., 2015).

To overcome this limitation, TAAC mixes a white noise with a noise generated from the stimulator-specific TMS *click* and tailored in real-time (in terms of time and frequency transformations) to match each individual’s *click* perception (See Supplementary methods). The noise obtained, named customized noise, is effective at safe (with respect to NIOSH and OSHA guidelines) and lower SPL with respect to the standard noises. Notably, this SPL reduction results in a major reduction (between 61% and 83% less) in terms of sound intensity. Thus, it can be safely employed even in case of long TMS-EEG sessions.

In case of patients unable to report their perception (e.g., disorders of consciousness, infants, newborns), TAAC can be associated with tools for the real-time visualization of evoked potentials (Casarotto et al., 2021) as in our electrophysiological proof-of-concept. This way, the experimenter can document in real time whether the AEP is present (typically characterized by the N100 and P200 components – Fig.1E, left side). Indeed, the documented absence of the AEP (Fig.1E, right side) is the most reliable criterion to verify the efficacy of the masking noise.

TAAC provides a major advancement in the reduction of AEP during TMS-EEG experiments, standardizing the generation of safe masking noises and effectively preventing the auditory contamination of TEPs by employing individualized procedures. TAAC is open-source and can potentially mask any transient sound, implying further improvements whether combined with technical advancements (e.g., quiet TMS Peterchev et al., 2015) and potentially extensive applications in any analogous setup.

## 5. Acknowledgements

We thank Brian L. Edlow for insightful discussion and comments on the manuscript.

