## Supplementary materials for "TAAC - TMS Adaptable Auditory Control: a universal tool to mask TMS click"

*TAAC - TMS Adaptable Auditory Control:*

*a universal tool to mask TMS click*

| **Noise** | **Type of noise** | ***Click* Distortion** | **White noise / *Click* noise ratio** | **Inter-*click*** | | |
| --- | --- | --- | --- | --- | --- | --- |
|  |  |  |  | **Fixed** | **Minimum** | **Maximum** |
| **White** | White | N/A | 100%  (only WBN) | N/A | N/A | N/A |
| **Adapted** | *Click*-based | N/A | 0%  (only CBN) | 1 | 1 | 50 |
| **Customized 1** | WBN + CBN | Broadband shuffling | 80% | 10 | 1 | 500 |
| **Customized 2** | WBN + CBN | Broadband shuffling | 80% | 100 | 1 | 10000 |

**Table S1**: Masking noise parameters

Parameters employed to generate each masking noise (N/A: Not applicable; Inter *click* latencies are reported in samples).

| **Subj** | **Real TMS** | | | | **Sham TMS** | | | |
| --- | --- | --- | --- | --- | --- | --- | --- | --- |
|  | **WN** | **AN** | **CN1** | **CN2** | **WN** | **AN** | **CN1** | **CN2** |
| **1** | 89.5 | 92.4 | 84.6 | 90.2 | 92.3 | 96.2 | 88.6 | 94.0 |
| **2** | 79.9 | 85.4 | 77.5 | 76.0 | 77.0 | 79.2 | 71.9 | 66.5 |
| **3** | 90.1 | 94.7 | 84.6 | 88.6 | 87.5 | 90.5 | 81.5 | 84.1 |
| **4** | 86.0 | 89.0 | 80.3 | 80.7 | 78.5 | 84.4 | 74.0 | 77.3 |
| **5** | 85.1 | 83.3 | 75.9 | 74.6 | 81.1 | 80.7 | 75.9 | 76.0 |
| **6** | 86.8 | 86.4 | 80.3 | 79.6 | 75.3 | 80.7 | 75.9 | 74.6 |
| **7** | 84.2 | 88.2 | 83.7 | 82.5 | 81.1 | 86.4 | 79.0 | 79.6 |
| **8** | 87.5 | 92.4 | 81.5 | 84.9 | 81.1 | 87.3 | 83.7 | 78.5 |
| **9** | 78.5 | 84.4 | 77.5 | 79.6 | 75.3 | 80.7 | 71.9 | 73.0 |
| **10** | 77.0 | 84.4 | 80.3 | 71.1 | 75.3 | 80.7 | 77.5 | 74.6 |
| **11** | 89.5 | 88.2 | 86.3 | 83.3 | 83.3 | 84.4 | 81.5 | 79.6 |
| **12** | 88.9 | 93.6 | 88.6 | 81.6 | 87.5 | 85.4 | 79.0 | 83.3 |
| **13** | 90.7 | 92.4 | 84.6 | 87.5 | 86.8 | 88.6 | 80.3 | 89.2 |
| **14** | 85.1 | 89.0 | 79.0 | 80.7 | 78.5 | 83.3 | 75.9 | 77.3 |
| **15** | 86.8 | 89.0 | 85.5 | 85.6 | 86.8 | 91.1 | 77.5 | 84.9 |
| **16** | 87.5 | 88.2 | 84.6 | 85.6 | 88.2 | 91.1 | 84.6 | 86.2 |
| **17** | 91.7 | 94.1 | 88.6 | 86.9 | 91.2 | 93.6 | 86.3 | 88.6 |
| **18** | 83.3 | 85.4 | 75.9 | 74.6 | 82.3 | 85.4 | 75.9 | 77.3 |
| **19** | 88.2 | 89.0 | 86.3 | 83.3 | 86.0 | 84.4 | 80.3 | 82.5 |
| **20** | 92.3 | 92.4 | 91.1 | 87.5 | 81.1 | 90.5 | 77.5 | 78.5 |

**Table S2**: Masking noise efficacy

Minimum SPL (dB) at which each noise was effective in masking the TMS click for each subject and condition

**Supplementary methods**

Subject-specific TAAC customized noise tailoring

While here we test the SPL required to mask TMS *click* through two pre-customized noises, TAAC allows generating fully customized masking noises strictly tailored on each and every subject. To do this, based on our experience, we recommend the following dichotomic questions to be posed to the subject and the respective modifications of the main features of the customized noise to be performed in the *Test noise* condition.

·	Do you perceive the high-pitched component of the TMS *click*?

1. Progressively increase the *Noise resampling* value

2. Progressively increase the *White noise resampling* value

3. Progressively increase the *White noise – click noise ratio* value

4. Progressively reduce the Inter-click values (e.g., rounded half values, all >1)

·	Do you perceive the low-pitched component of the TMS *click*?

1. Progressively reduce the *White noise resampling* value

2. Progressively reduce the *White noise – click noise ratio* value

3. Progressively reduce the *Noise resampling* value

4. Progressively increase the Inter-click values (e.g., doubled values, all >1)

To customize the noise on a single subject, we suggest starting the parameterization from the default TAAC parameters, as following:

·	*Noise resampling*: 96000 Hz

·	*White Noise resampling*: 96000 Hz

·	*Noise volume*: 100%

·	*White noise – Click noise ratio*: 80%

·	*Fixed inter-click (samples)*: 10

·	*Min inter-click (samples)*: 1

·	*Max inter-click (samples)*: 500

·	*Maximum volume reduction (%)*: 0

·	*Left max shift (samples)*: 0

·	*Right max shift (samples)*: 0

Preprocessing of electrophysiological data

We acquired EEG data through a 62-channel cap (EasyCap) connected to a TMS-compatible BrainAmp DC amplifier (BrainProducts GmbH, Munich, Germany). EEG signal was recorded with a resolution of 0.5 µV at DC with a high-frequency cutoff of 1 kHz and digitized at a sampling rate of 5 kHz.

Data analysis was performed using Matlab R2016b (The MathWorks) based on custom-made scripts (Fecchio et al., 2017). Artifact-contaminated epochs were manually rejected by visual inspection. The TMS step-artifact was removed using a linear interpolation between -2 and 5 ms around the TMS pulse. Then, EEG data were detrended, high-pass filtered (1 Hz, Butterworth, 3rd order), epoched from -600 to 600 ms around the TMS pulse, and referenced to the average reference. Independent component analysis (ICA, EEGLAB *runica* function) was applied to remove eye blinks/movements and spontaneous scalp muscle activations. Finally, the data were low-pass filtered (45 Hz, Butterworth, 3rd order) and downsampled at 1 kHz. No bad channels were identified, and 109 good epochs were averaged to obtain the potential evoked by TMS in each condition.
